# Instrument-free quantitative detection of analytes using a novel combo lateral flow assay

**DOI:** 10.1101/2025.09.04.674023

**Authors:** Xiaolin Sheng, Navya Saxena, Xiaofeng Xia

**Affiliations:** Instanosis Inc., 1004 W 9^th^ Avenue, King of Prussia, PA 19406

## Abstract

Lateral flow assays (LFA) are widely regarded as the cheapest, fastest, and easiest point-of-care (PoC) tests available.^1,2^ A key advantage is that results can be visually interpreted without the need for specialized equipment, enabling on-site applications while keeping costs low.^3^ However, conventional LFAs are typically qualitative, limiting their use to simply detect the presence or absence of analytes.^4^ Additional limitations include narrow dynamic ranges and false negatives due to the Hook Effect in sandwich LFAs.^5^

In this work, we present a combo lateral flow assay (CLFA) that integrates both sandwich and competitive assay formats, enabling the quantitative detection of analytes across a broad concentration range without an external instrument. The precision of CLFA measurements can be further enhanced with artificial intelligence (AI)-based image analysis via a smartphone app. Importantly, CLFA is not subject to the Hook Effect.

CLFA is easily implemented by adding just one extra test line to the standard LFA design, resulting in minimal additional manufacturing effort and cost. Given these benefits, we anticipate that CLFA will have broad applications in diagnostics and research applications. Several examples are provided to illustrate its potential.

## Introduction

Lateral flow assays (LFAs) are among the most widely used diagnostic tests.^2,6,7^ Their broad public acceptance stems first from their long-standing use in pregnancy testing,^8^ and more recently from their widespread adoption during the COVID-19 pandemic.^9,10^ It is estimated that over two billion LFA devices are produced each year.^8^

The simplicity and low cost of LFAs have made them a popular point-of-care (PoC) tool across diverse fields, including research and development, clinical diagnostics,^11^ environmental monitoring,^12^ and safety assessments^13,14^. Additionally, LFAs offer practical advantages such as long shelf life and storage without refrigeration, making them particularly suitable for use in developing countries, small clinics, remote areas, and even in battlefield conditions.^3^

LFAs typically rely on one of two main sensing strategies: sandwich assays or competitive assays. The sandwich assay^15^ is likely the most commonly used approach for detecting mid-to large-sized analytes (>1 kDa), such as biomarker proteins, antibodies, and pathogens. In this format, the target molecule is captured between a detection bioreceptor and a test-line capture bioreceptor, generating a signal that increases proportionally with the concentration of the target in the sample.^16^ Competitive assays^17^ come in two main formats. In the first, the target in the sample competes with a labeled target (or an alternative molecule with lower affinity for the bioreceptor) from the conjugate pad for binding to the test-line capture bioreceptor. In the second format, the target in the sample competes with immobilized target molecules on the test line for binding to the labeled bioreceptor.^18^

Despite its advantages, LFA remains primarily a qualitative technology. Quantitative measurements typically require a dedicated reader device.^19^ Even with a reader, however, the assay’s dynamic range is limited, usually spanning no more than two orders of magnitude. This limitation is more pronounced in competitive LFAs, which generally have lower sensitivity and a narrower dynamic range compared to sandwich LFAs.^20^ Additionally, sandwich LFAs are susceptible to the Hook Effect,^5^ where excessively high analyte concentrations can cause the test line to disappear, leading to false-negative results. To avoid this, serial dilutions of samples are often necessary, which increases both the workload and the complexity of the assay.

Here, we present a combo lateral flow assay (CLFA) that combines both sandwich and competitive assay formats to enable quantitative analysis without the need for instrumentation, while completely eliminating false negatives caused by the Hook Effect. Our design leverages the distinct response ranges of sandwich and competitive assays to expand the dynamic range and allow quantification of analytes by visual inspection. This novel approach transforms LFA interpretation from a simple color intensity measurement into a pattern recognition task. This not only makes visual assessment easier but also enhances compatibility with modern image processing and machine learning algorithms. To showcase the versatility of CLFA, we demonstrated its applications in both the development and manufacturing of therapeutic biologics, as well as in PoC diagnostics.

## Methods

### Materials

The following materials were used in producing the CLFA strips: Lateral flow backing cards (Millipore Sigma, Catalog HF135MC100); CN95 nitrocellulose membrane (Satorius, Catalog 1UN95ER100025NT); Glass fiber conjugation pad (Millipore Sigma, Catalog GFCP103000); CF6 absorbent pad paper (Cytiva, Catalog 8116-2250); Standard 17 glass fiber membrane (Cytiva, Catalog 8134-2250).

The following materials were used as standards in CLFA testing: Human IgG1 (Abcam, Catalog AB90283); Fresh human whole blood (HumanCells Biosciences, Catalog FP-004-10). Premade AAV2 virus reference (AAVnerGene, Catalog DE000000-AAV2).

### CRP antibody development

Native human CRP was purchased from Biorad (Hercules, Catalog PHP277) and used as immunogen. Rabbit immunization was performed by Cocalico Biologicals (Stevens, PA) following a standard procedure.^21^ All animal maintenance, care and use procedures were reviewed and approved by the company’s Institutional Animal Care and Use Committee (IACUC). Antibodies were developed and cloned using a single B cell method to obtain the coding sequences.^22^ Recombinant antibodies were then expressed and purified from Expi293F cells.

### Cell transfection

Expi293F cells (Thermofisher, Catalog A14527) were transfected with the expression vector using the ExpiFectamine 293 transfection reagent (Thermofisher, Catalog A14525) following the product instruction. To establish stable expression cell lines, a GFP expression cassette was added to the expression vector, and the single transfected cells expressing GFP were sorted with a BD FACSMedody (BD Biosciences) into 96-well plates. The cells were cultured for 2 weeks and transferred to 6-well plates for further expansion of 2 weeks. After that the expression levels were measured using CLFA and ELISA.

### Gold nanoparticle preparation and conjugation

Gold nanoparticles with an average diameter of 40 nm were synthesized using the Turkevich Method.^23,24^ Briefly, 100 mL of 1 mM HAuCl4 solution was boiled under stirring. Ten ml of 38.8 mM sodium citrate was quickly added to the boiling HAuCl_4_ solution while the stirring was kept for 15 min. After that the solution was cooled down to room temperature. The size of the nanoparticle was confirmed using UV–visible spectroscopy, with the peak at 530nm.^25^

The gold nanoparticles were functionalized with antibodies by physical adsorption.^17^ Briefly, the pH of the AuNP solution was first adjusted to 9.0 using 100 mM potassium carbonate solution. Then, 40 μg antibodies were mixed with 10 mL of gold nanoparticle overnight at 4°C. A solution of 1% BSA was further used to block the unreacted sites on the surface of the AuNPs for 2 h at 4°C. Finally, the functionalized gold nanoparticles were collected by centrifugation at 4,000 g for 20 min, and then re-suspended in 2.5 mL of PBS, pH 7.4, containing 0.1% BSA. The following antibodies were conjugated in this study: mouse anti human IgG Fc (Thermofisher Catalog MA1-30178) For human IgG Fc CLFA; rabbit anti CRP clone 194 for CRP CLFA; rabbit anti AAV2 antibody clone 15G4 (Genscript, Catalog A02202) for AAV2 CLFA.

### Strip fabrication

For human IgG Fc CLFA, the control line was prepared by coating 1 mg/mL rabbit anti mouse IgG Fc polyclonal (Jackson Immuno Research, Catalog 315-005-046); The sandwich test line was prepared by coating 0.5 mg/mL mouse anti human IgG Fc (Thermofisher, Catalog MA5-16858); The competitive test line was prepared by coating 0.5 mg/mL Human IgG (Innovative Research, Catalog IHUIGGAP10MG).

For AAV2 virus particle CLFA, the control line was prepared by coating 1 mg/mL protein G (Prospec, Catalog PRO-402); The sandwich test line was prepared by coating 0.5 mg/mL anti AAV2 antibody clone 15G4 (Genscript, Catalog A02202); The competitive test line was prepared by coating 1 mg/mL recombinant AAV2 capsid protein VP3 (Fisher Scientific, Catalog 50-253-3991).

For CRP CLFA, the control line was prepared by coating 1 mg/mL protein G; The sandwich test line was prepared by coating 0.25 mg/mL rabbit anti CRP clone 35; The competitive test line was prepared by coating 0.5 mg/mL native human CRP protein (Biorad, Catalog PHP277).

### ELISA

Human IgG concentration was measured using an IgG quantification ABC kit (Kerafast, Catalog ABCK0001). CRP concentration was measured using a human CRP ELISA kit (Millipore Sigma, Catalog RAB0096).

### AI-based image analysis

A combination of a classifier at anchor points to create buckets and a regressor for each bucket was used. The modeling framework begins by extracting a compact, 23-dimensional feature vector from each lateral flow strip image. After cropping out a central region and converting it to both grayscale and HSV (Hue, Saturation, Value), the pipeline applies a black hat morphological transform to accentuate faint bands, then computes smoothed intensity profiles along the vertical axis. Three peak positions (control line and the two test lines) are located based on prominence, and for each band the algorithm measures peak height, integrated area, width, local grayscale and raw V (Value) □hannel statistics. Two additional features capture the ratio of the first and second test line response, giving a rich representation of both absolute and relative signal intensity.

Next, a Random Forest classifier serves as a gate that assigns each strip to one of six concentration buckets. This Random Forest classifier has 200 Decision Trees. Each tree in the ensemble chooses whichever feature and threshold maximizes the information gain at that node, and the final bucket assignment is the majority vote across all 200 trees. An example is provided in **Supplemental Figure 1**).

Following bucket assignment, each strip’s feature vector is passed to a dedicated regression pipeline. First, the 23 features are standardized using a StandardScaler fitted on that bucket’s training samples. The scaled vector is then input to a histogram gradient boosting regressor configured with five hundred boosting iterations, a learning rate of 0.05, a maximum tree depth of eight, and an absolute error loss function. For the three lowest non-zero buckets (0.1-1, 1-10, and 10-100 µg/mL), the raw regressor output is subsequently adjusted by an isotonic regression model with out of bounds values clipped to correct systematic bias and residual nonlinearity.

Finally, the calibrated concentration estimate is given. Strips lacking a detectable control line bypass regression and are flagged invalid. This bucket specific mixture of experts enables the pipeline to tailor modeling complexity to each concentration interval while maintaining efficient on-device inference.

The image dataset included over 5000 smartphone captured IgG lateral flow strips at 19 nominal concentrations (0, 0.1, 0.5, 1, 5, 7, 10, 30, 50, 70□µg/mL and 0.1, 0.3, 0.5, 0.7, 1, 3, 5, 7, 10□mg/mL). All images enumerated per concentration and, using a fixed random seed, each bucket was split into 80% for training and 20% for held out testing to ensure balanced representation across the full dynamic range. The held-out test was captured under varied lighting and slight strip tilt to rigorously evaluate generalization.

## Results

### CLFA design

CLFA is implemented by combining sandwich and competitive LFA formats, as illustrated in **Figure 1**. The strip is consisted of a standard sample pad for sample application, followed by a conjugate pad containing gold nanoparticles (GNPs) conjugated with analyte detection antibodies [5][6]. As the sample migrates onto the nitrocellulose membrane, it first encounters the sandwich test line, which is coated with an analyte capture antibody to retain GNPs bound to the analyte. The sample then continues to the competitive test line, coated with the analyte itself, which captures GNPs that are unbound to analytes. Any remaining GNPs continue migrating to the control line, which is coated with a secondary antibody or protein G to confirm proper flow. Because the sandwich and competitive test lines respond inversely to analyte concentration — with the competitive assay typically being less sensitive than the sandwich assay — increasing analyte levels first lead to the appearance and intensifying of the sandwich test line, then the diminishing and eventually the elimination of the competitive test line. This generates a distinct line pattern across a much broader dynamic range.

**Figure 1.**
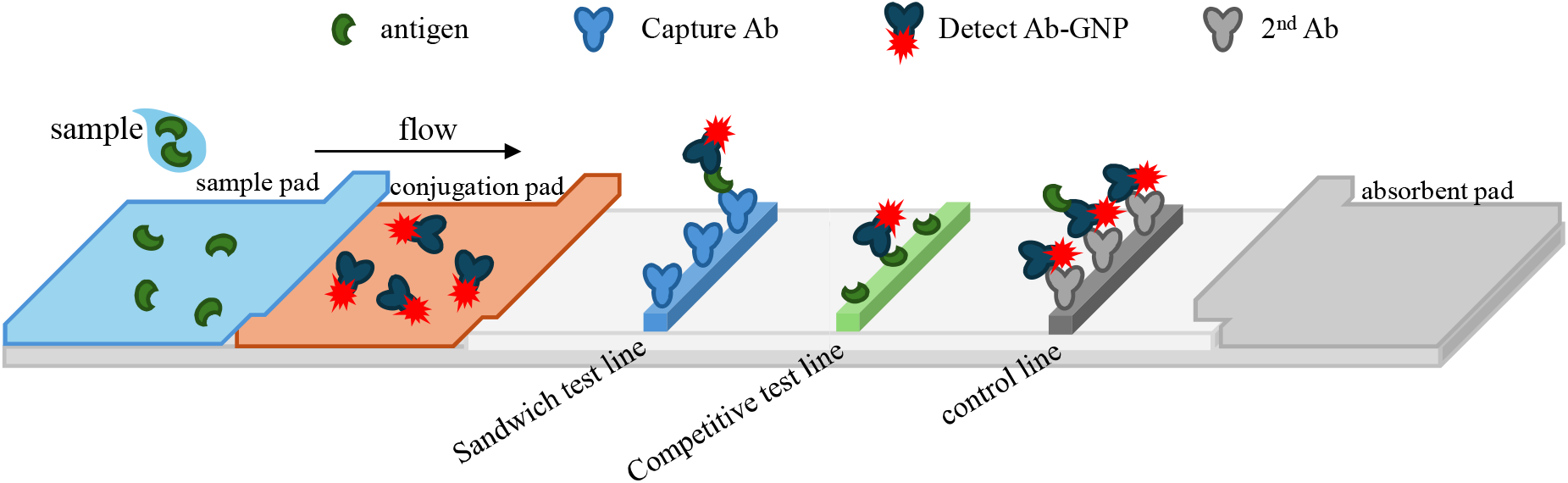
Schematic illustration of CLFA. CLFA is constructed by combining sandwich and competitive LFA formats, incorporating a sandwich test lines, a competitive test line and a control line. Due to the opposite responses of the sandwich and competitive test lines, along with their differing dynamic ranges, CLFA produces distinct line patterns across a broad concentration range that are easily recognizable by the naked eye. Compared to traditional LFA, the CLFA line patterns are also better suited for modern AI-based image analysis.

By interpreting the combined line pattern instead of relying solely on the color intensity of a single line, CLFA greatly simplifies visual assessment. The dual-line information enhances both accuracy and dynamic range when analyzed via either naked eyes or imaging technologies.

Furthermore, CLFA inherently prevents inaccurate or false-negative results from the Hook Effect [3]: true negative samples display both the control and competitive test lines, whereas in the case of the Hook Effect with the presence of excessively high analyte concentrations, only the control line appears.

### CLFA application in recombinant protein and antibody production

To demonstrate the application of CLFA, we developed a test strip for detecting the Fc region of human IgG, as described in the Methods. Standard samples containing human IgG1 concentrations ranging from 0 µg/mL to 10 mg/mL were tested, and the results are shown in **Figure 2**. Distinct line patterns were observed across this 5-log concentration range. In negative samples (0 µg/mL IgG1), two lines appeared: the control line (C) and the competitive test line (T1), with T1 more intense than C. At 0.1 µg/mL IgG1, the sandwich test line (T2) began to appear and increased in intensity with higher IgG1 concentrations, reaching parity with T1 at 10 µg/mL. As IgG1 levels increased, T1 intensity decreased while T2 continued to strengthen. At 100 µg/mL, T1 was faint and T2 was roughly equal in intensity to C. When the concentration reached 1 mg/mL, T1 disappeared, and T2 weakened to below C intensity. Beyond 10 mg/mL, both T1 and T2 vanished, leaving only the control line visible. Throughout this concentration range, the presence, absence, and relative intensities of C, T1, and T2 produced distinct patterns that can be visually recognized, enabling semi-quantitative estimation of human IgG1 levels without the need for instrumentation.

**Figure 2.**
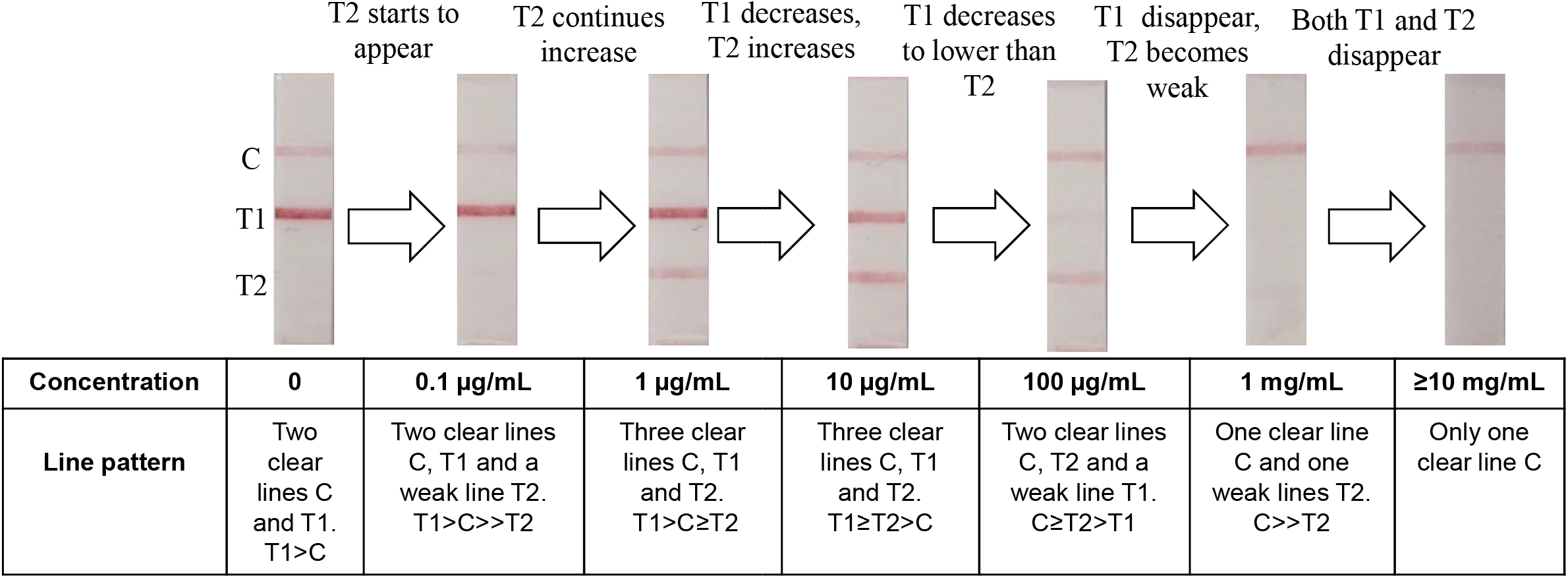
Human IgG CLFA. Human IgG1 standard samples were prepared in PBS buffer and tested using the CLFA strip. The resulting line patterns, spanning human IgG1 concentrations across five orders of magnitude, are shown. Each concentration produces a distinct line pattern that can be easily distinguished by the naked eye, enabling semi-quantitative assessment without instrumentation.

To demonstrate the application of the human IgG Fc CLFA, we transfected 293 cells with a construct expressing a human IgG1 Fc-tagged PD1 protein. One hundred µL samples were collected daily over 5 days and tested using both CLFA and ELISA (**Figure 3A**). On Day 1, a faint T2 line was visible, corresponding to a concentration between 0.1 and 1 µg/mL based on the standards in **Figure 2**, consistent with the ELISA result of 0.8 µg/mL. By Day 2, T2 became clearer but remained weaker than T1, indicating a concentration of 1–10 µg/mL, while ELISA measured 2.6 µg/mL. On Day 4, C, T1, and T2 lines showed similar intensity, suggesting a concentration of 10–100 µg/mL, a pattern that persisted on Day 5. ELISA confirmed concentrations of 32.5 µg/mL and 37.2 µg/mL on Days 4 and 5, respectively, aligning with the CLFA results. Since the CLFA assay requires only 5 minutes with minimal handling, this demonstrates its utility for convenient and rapid monitoring of recombinant protein expression post-transfection.

**Figure 3.**
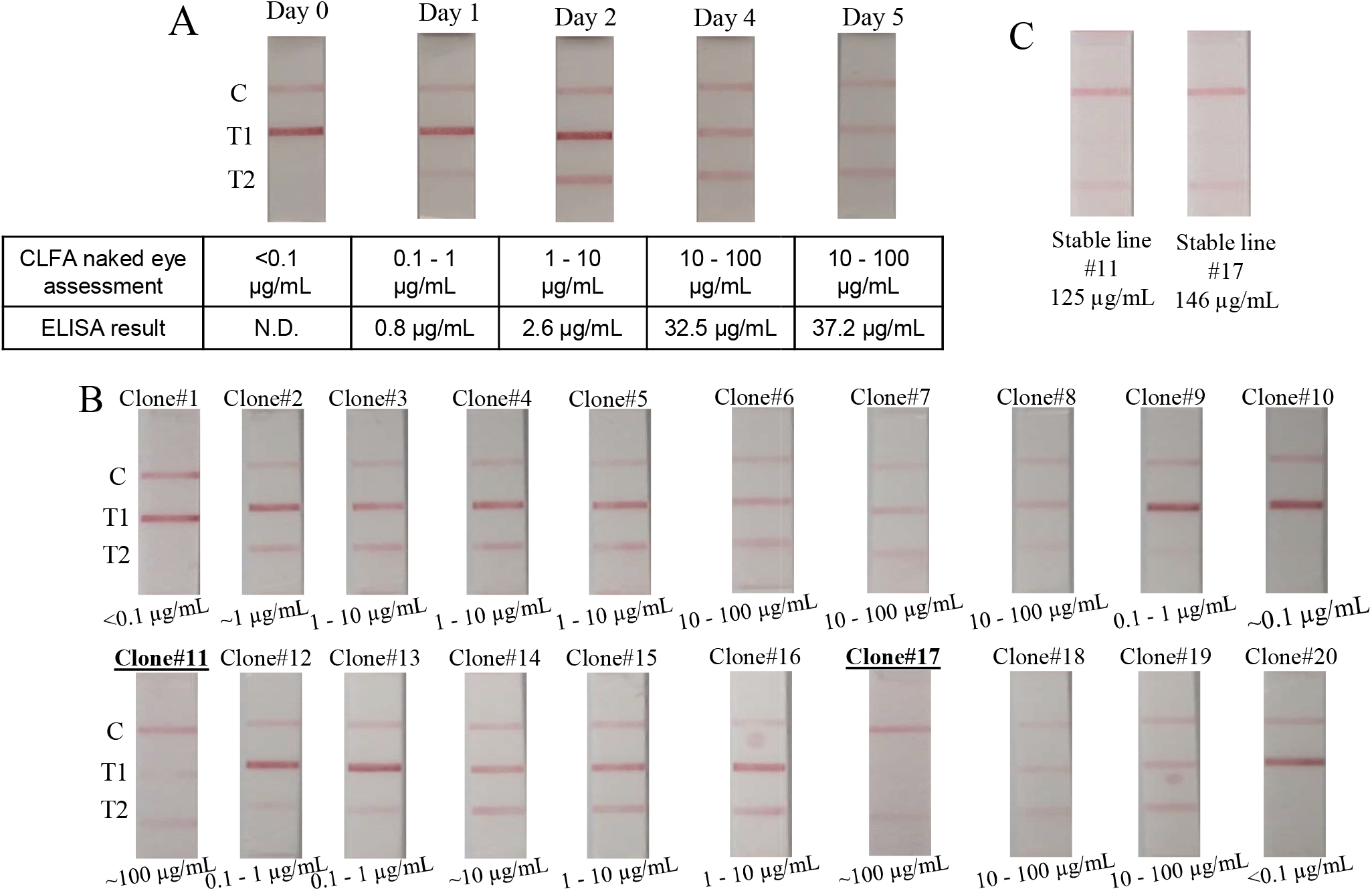
Examples of CLFA applications in recombinant protein production and antibody stable cell line development. **(A)** Monitoring transient expression of recombinant Fc-tagged PD1 protein using CLFA. Samples were collected daily, and protein concentrations were assessed by both CLFA (via naked eye observation) and ELISA. CLFA revealed peak expression after Day 4, which was confirmed by ELISA. **(B)** Clone selection for stable Herceptin-expressing 293 cell lines using CLFA. Twenty clones were screened, and two clones (bold and underlined) were selected based on CLFA line patterns. **(C)** ELISA validation of antibody expression levels in the selected clones, confirming high Herceptin production.

We further evaluated CLFA in recombinant antibody production by transfecting 293 cells with a construct expressing the anti-HER2 antibody Herceptin, co-expressed with GFP for cell sorting. After 4 weeks, when transient expression subsided, we isolated 20 single-cell clones using FACS. The single cells were cultured and assessed for Herceptin expression levels using CLFA. The resulting line patterns enabled semi-quantitative evaluation, allowing us to identify clones #11 and #17 as top producers (**Figure 3B**). These clones were expanded into stable cell lines, and ELISA confirmed Herceptin expression levels of 125 µg/mL and 146 µg/mL, respectively, consistent with the CLFA assessments (**Figure 3C**). This study demonstrates that CLFA can support high-throughput clone selection with rapid, simple procedures, enhancing both the speed and reliability of stable antibody-expressing cell line development, which is critical for pharmaceutical production.

### CLFA application in virus titering

Adeno-associated virus (AAV) has become the predominant gene delivery tool in gene therapy in recent years [17]. Accurate AAV titration is critical for therapeutic development, manufacturing, and quality control. A rapid, low-cost, and simple titration method could significantly improve screening throughput and enable closer monitoring throughout both development and production processes.

To explore the use of CLFA for AAV titration, we developed a CLFA assay targeting AAV2, with the competitive test line coated with AAV2 capsid VP3 protein and the sandwich test line coated with an anti-AAV2 antibody. Standard samples ranging from 10□ to 10^13^ genome copies (GC)/mL were prepared in DMEM supplemented with 10% FBS, alongside a blank control. As shown in **Figure 4**, distinct line patterns were observed across this 4-log titer range. In the negative control, only the control (C) line and the competitive test line (T1) appeared. At 10□GC/mL, the sandwich test line (T2) became faintly visible, and its intensity increased steadily, becoming clear at 10^1^□GC/mL. As T2 intensity increased, T1 intensity decreased, becoming weaker than T2 at 10^11^ GC/mL. This trend continued, with T2 fading at 10^12^ GC/mL and disappearing entirely at 10^13^ GC/mL.

**Figure 4.**
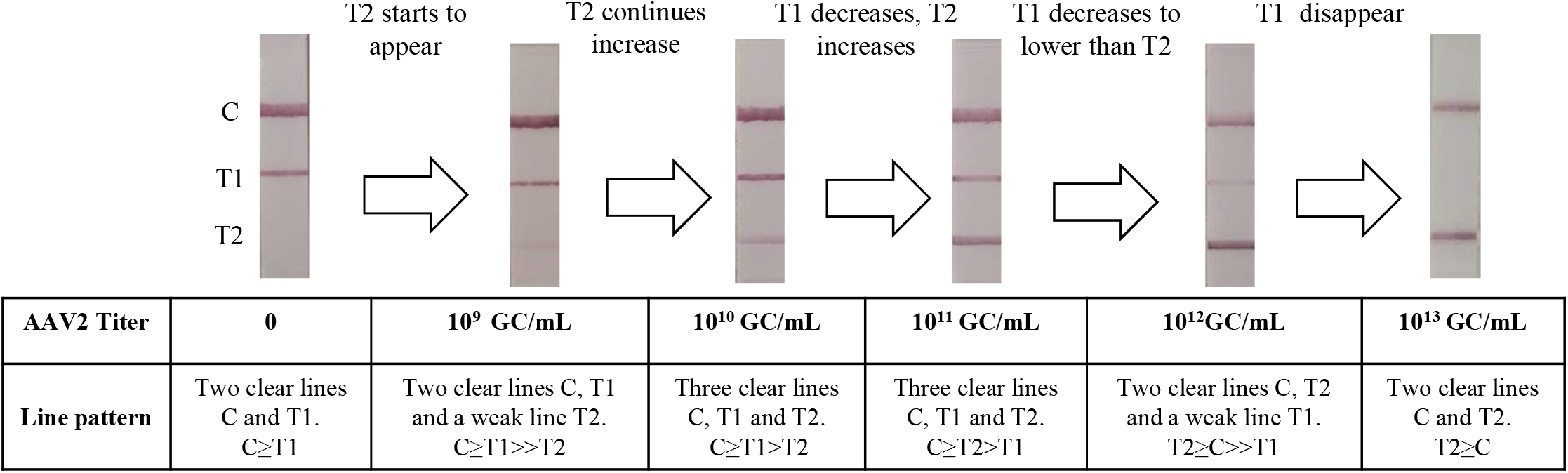
AAV2 CLFA. AAV2 virus particle standard samples were prepared in DMEM with 10% FBS and tested using CLFA. The resulting line patterns, spanning AAV2 titers across four orders of magnitude, are shown. These patterns exhibit distinct features that can be easily recognized by the naked eye for rapid, semi-quantitative assessment.

These easily distinguishable line patterns enable rapid, cost-effective, and instrument-free assessment of AAV titers, providing a practical tool for monitoring virus production in both research and manufacturing settings.

### CLFA application in point-of-care (POC) diagnosis

Lateral flow assays (LFAs) are widely used in diagnostics due to their simplicity, making them ideal for PoC applications. However, without an instrument reader, LFAs usually only provide qualitative results and have a narrow dynamic range, as line intensity typically reaches its maximum within about one order of magnitude. To showcase the advantages of the novel CLFA assay for PoC diagnostics, we developed an assay to quantitatively measure C-reactive protein (CRP) levels in human blood samples. CRP is an acute-phase reactant protein used both as a cardiac disease and a nonspecific inflammation marker, with distinct concentrations for each application.^26^ Clinically, levels below 0.3 mg/dL indicate average risk while those from 0.3 to 1.0 mg/dL predict high risk for coronary heart diseases. Levels from 1.0 to 10.0 mg/dL represent elevation often linked to systemic inflammation or autoimmune diseases; levels above 10.0 mg/dL indicate marked elevation, usually associated with acute bacterial or viral infections or major trauma. Severe elevations exceeding 50.0 mg/dL typically result from acute bacterial infections.

To evaluate whether the CLFA could differentiate blood samples across these CRP ranges, we developed a PoC kit comprising a sample loop which is frequently used for capillary blood collection in PoC applications, a sample vial, a dropper, and a cassette containing the CLFA strip (**Figure 5A**). The assay procedure (**Figure 5B**) involved collecting 10 µL of blood using the loop, diluting it 100-fold in 1 mL of PBS buffer within the sample vial, then applying three drops of the diluted sample onto the cassette via the dropper. After 5 minutes, results were read to determine CRP concentration. For standard preparation, fresh human whole blood was first measured by ELISA to have a CRP level of 0.2 mg/dL. Higher concentration samples were prepared by spiking additional CRP protein. A blank control sample was created by removing CRP using beads coated with CRP antibodies and confirmed by ELISA. When tested with the CLFA kit, distinct line patterns emerged that corresponded to clinically relevant CRP ranges (**Figure 5C**). Normal samples with CRP below 0.3 mg/dL showed two clear lines—C and T1—while the T2 line was absent or very faint. Minor CRP elevation (around 0.3 mg/dL) produced a visible T2 line with intensity lower than T1. As CRP increased, T1’s intensity rose, equaling T2 at 1 mg/dL, indicating moderate elevation. In the marked elevation range (10 to 50 mg/dL), T2 became stronger than T1, which weakened. At severe elevations (>50 mg/dL), the T1 line disappeared, leaving only the C and T2 lines visible. These distinct and easily recognizable line patterns align well with clinically significant CRP levels, providing a simple, user-friendly method for laypersons to monitor CRP at home.

**Figure 5.**
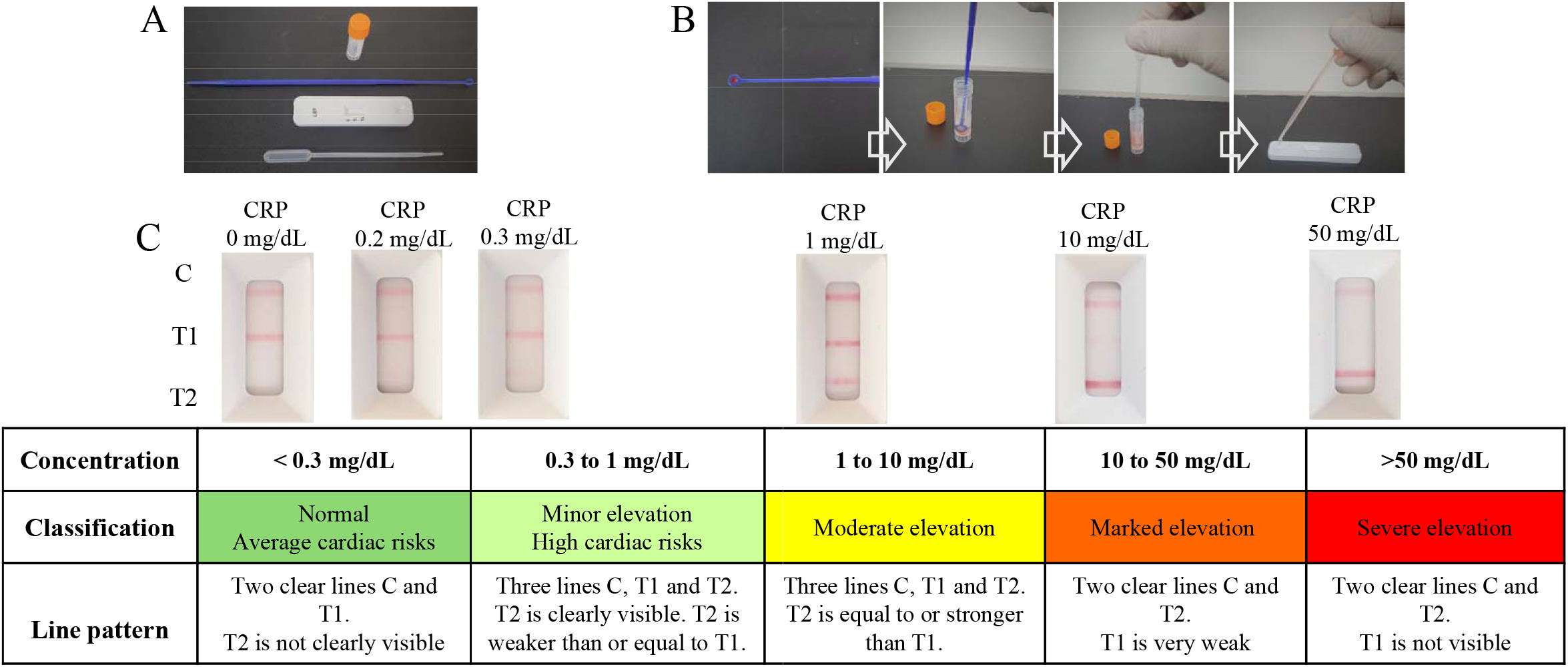
A CLFA PoC kit for CRP testing in human whole blood. **(A)** Kit components include a sample vial containing 1 mL PBS buffer, a loop for collecting 10 µL of whole blood, a cassette housing the test strip, and a dropper for applying the sample onto the cassette. **(B)** Assay procedure: 10 µL of whole blood is collected using the loop and diluted with 1 mL of PBS buffer in the vial. After mixing thoroughly, three drops of the diluted sample are applied onto the cassette. Results are read 5 minutes later. **(C)** CLFA results: Line patterns corresponding to samples with CRP concentrations at clinical cutoff values are shown. Clinical classifications can be easily identified by visual inspection of the CLFA line patterns.

### AI enhanced CLFA quantitative measurement

CLFA turned analyte measurements from simple line intensity readings to line pattern recognition, providing a richer source of information. Naked eye inspection can already assess the semiquantitative analyte concentration across a broad range. We hypothesize that more accurate and quantitative measurements can be achieved through image analysis on a cellphone, especially with the recent rapid advancement of AI image recognition technology. Towards this goal, we developed a pipeline as described in Methods that combines peak based feature extraction (**Supplemental Figure 2**) with a mixture of experts scheme to deliver true concentration regression from 0 to 10,000 µg/mL. Each strip is first routed through a balanced random forest classifier that assigns it to one of six predefined concentration buckets. Within each bucket a StandardScaler normalized histogram gradient boosted regressor predicts the analyte level, and isotonic regression calibration corrects residual nonlinearity in the lowest ranges.

The pipeline was evaluated using two key metrics: mean absolute error (MAE), which reports error in the same units as the assay (µg/mL), and the coefficient of determination (R^2^), which quantifies the fraction of concentration variance explained by the model.

A cellphone app was developed to guide image capture and perform concentration calculation using this pipeline (**Figure 6A**). On a held-out test set spanning the full range (0–10,000 µg/mL), the model achieved an R^2^ of 0.9844. The correlation between AI-predicted and actual concentrations, along with MAE values for each range, is shown in **Figure 6B**.

**Figure 6.**
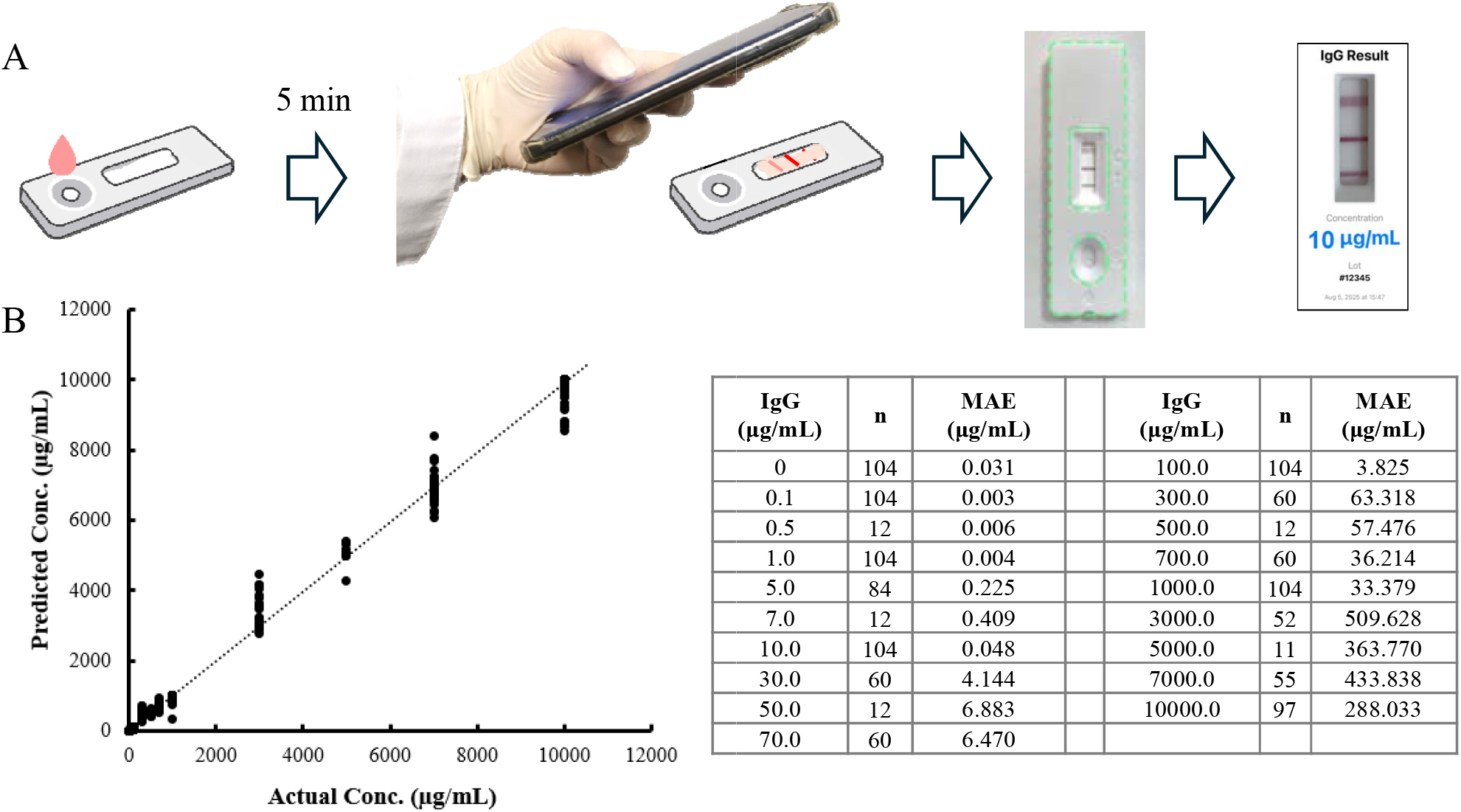
CLFA results read using a cellphone app with AI-assisted image analysis. **(A)** Workflow and the example user interface of the CLFA system operated via a cellphone app. **(B)** Accuracy evaluation of the results. Correlation between AI-predicted (dots) and actual concentrations (line) is shown on the left, and MAEs at individual concentrations are shown on the right. n, number of samples tested.

## Discussion

In this study, we present a simple yet highly effective approach to significantly enhance the functionality of traditional LFA technology. Given the proven ease of use and cost-effectiveness of conventional LFAs, the introduction of CLFA technology expands the scope of applications across research, diagnostics, therapeutic development, and manufacturing, as demonstrated by the examples in this paper.

CLFA is implemented by simply adding an additional test line to the standard LFA format. Because it uses the same conjugates and core components, this approach can be readily integrated into existing products and enables the development of new assays with minimal additional cost.

The central innovation of this method lies in incorporating both a sandwich test line and a competitive test line, which respond inversely to analyte concentration. This configuration generates line intensity patterns that are concentration-dependent, effectively adding a new dimension of data to the readout. The dual-response mechanism significantly improves assay readability and accuracy while expanding the dynamic range, as the sandwich and competitive formats respond in different concentration ranges. Although this study did not explore additional line configurations, it is plausible that increasing the number of lines, e.g., through variation in reagent concentration or affinity, could further enrich the signal pattern and enhance quantitative precision. Similarly, while traditional GNPs were used in this study, incorporating nanoparticles with different optical properties, such as highly sensitive gold nanorods^27^ or colored latex beads^10,28^, could also enhance pattern complexity and broaden assay functionality.

As demonstrated, these more intricate line patterns are well-suited for interpretation using modern AI technology, enabling the development of smartphone-based APPs capable of rapid, fully quantitative analysis even in resource-limited settings.^29–31^

This study focused on the instrument-free application of CLFA, with results interpreted by the naked eye. However, portable LFA readers suitable for PoC settings already exist.^32,33^ When paired with such readers, CLFA is expected to offer even greater precision and a broader dynamic range. Furthermore, the use of advanced nanoparticles—such as quantum dots,^34^ upconverting nanoparticles (UCNPs),^35^ or magnetic nanoparticles^36^ —could generate highly sensitive fluorescent or magnetic signals, pushing the accuracy and dynamic range of CLFA to unprecedented levels while preserving its low cost and rapid turnaround, making it a powerful tool for PoC diagnostics.

Beyond diagnostics, we also demonstrated the utility of CLFA in the research, development, and manufacturing of therapeutic biologicals. In these settings, CLFA offers a rapid, user-friendly, and relatively high-throughput screening method. Its broad dynamic range and resistance to the Hook Effect are particularly valuable, as concentrations of biologicals can range from a few µg/mL to over 10 mg/mL. In manufacturing especially, analyte levels often exceed the threshold where the Hook Effect occurs, leading to inaccurate measurements or false negatives with conventional sandwich ELISA based assays. CLFA effectively overcomes this limitation. In this study, we showed that analyte concentrations could be accurately assessed by visual inspection alone. Looking forward, integrating CLFA with portable readers could provide an even faster, more cost-efficient, higher throughput and easy-to-use alternative to current analytical methods, with the potential to become a standard tool in both research and industrial workflows.

## SUPPLEMENTARY FIGURES

**Supplemental Figure 1.**
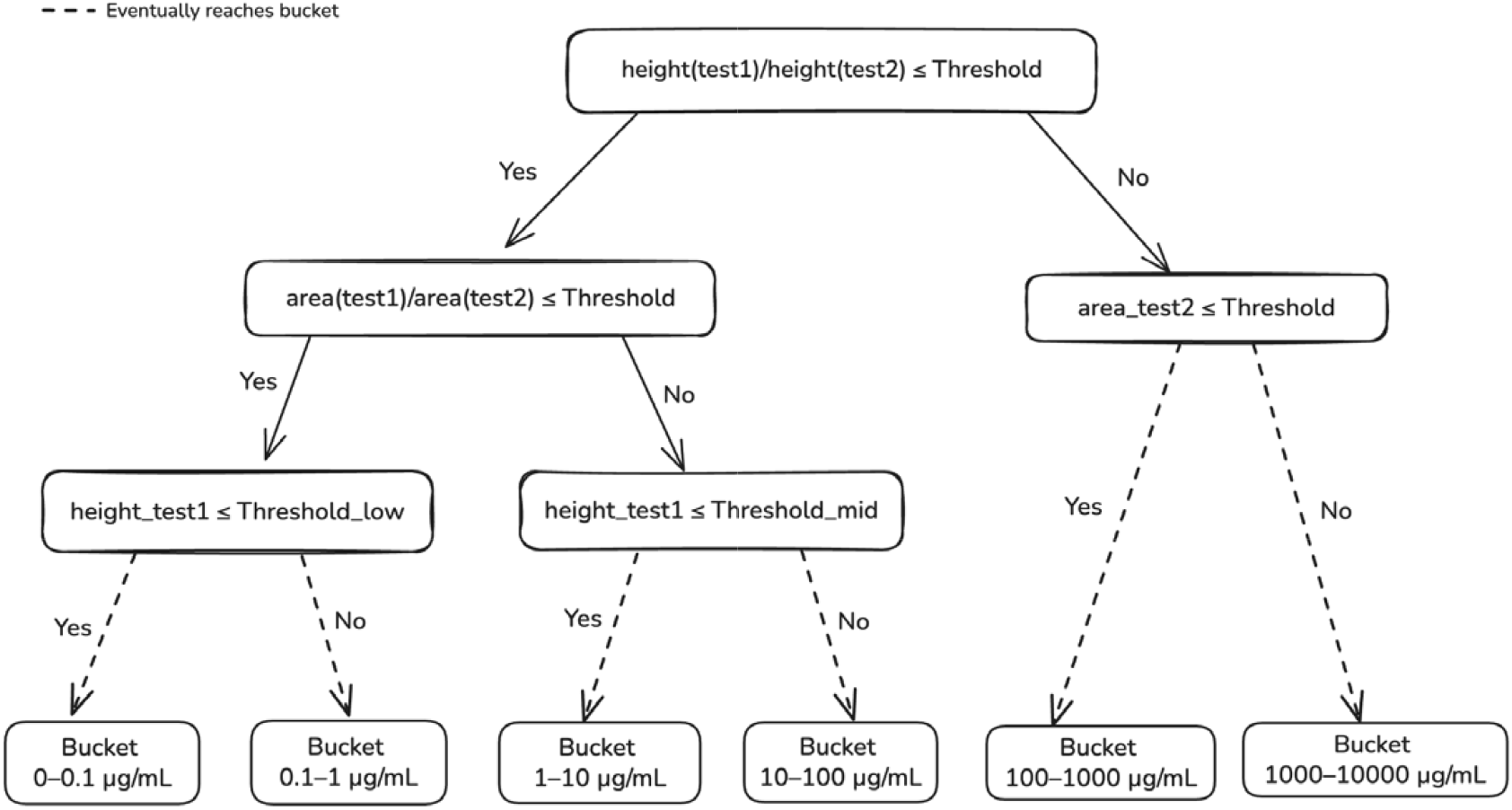
Example Decision Tree from the Random Forest Bucket Classifier.

**Supplemental Figure 2.**
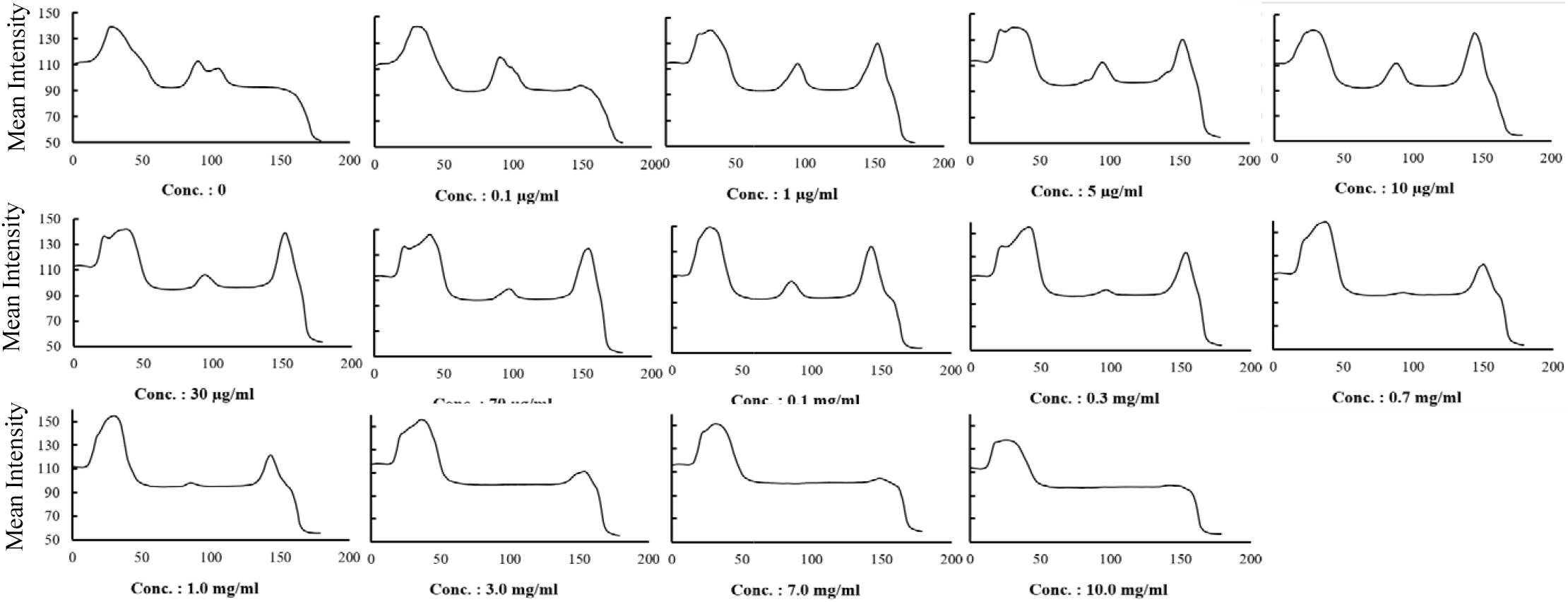
Signal intensity profile of the LFA strip. The x-axis shows scan position, and the y-axis shows measured intensity. Test line strength was obtained by subtracting background from the peak signal, and analyte concentrations were calculated from a standard calibration curve.

